# Leptin induces cell migration and invasion in a FAK-Src- dependent manner in breast cancer cells

**DOI:** 10.1101/631143

**Authors:** Juan C. Juárez-Cruz, Miriam Daniela Zuñiga-Eulogio, Monserrat Olea-Flores, Eduardo Castañeda-Saucedo, Miguel Ángel Mendoza-Catalán, Carlos Ortuño-Pineda, Ma. Elena Moreno-Godínez, Sócrates Villegas-Comonfort, Teresita Padilla-Benavides, Napoleón Navarro-Tito

## Abstract

Breast cancer is the most common invasive neoplasia, and the second leading cause of death associated with cancer in women worldwide. Mammary tumorigenesis is severely linked to obesity, the potential connection is leptin. Leptin is a hormone secreted by adipocytes, which contributes to the progression of breast cancer. Cell migration, metalloproteases secretion, and invasion are cellular processes associated with various stages of metastasis. These processes are regulated by the kinases FAK and Src. In this study, we utilized the breast cancer cell lines MCF7 and MDA-MB-231 to determine the effect of leptin on FAK and Src kinases activation, cell migration, metalloproteases secretion, and invasion. By Western blot we found that leptin activates FAK and Src, and induces the localization of FAK to the focal adhesions. Specific inhibitors of FAK and Src showed that the effect exerted by leptin in cell migration, and invasion in breast cancer cells is dependent on these kinases. Moreover, by gelatin zymmography we established that leptin promotes the secretion of the extracellular matrix remodelers, MMP-2 and MMP-9, in a FAK and Src dependent manner. Our findings strongly suggest that leptin promotes the development of a more aggressive invasive phenotype in mammary cancer cells.

## INTRODUCTION

Breast cancer is the most common invasive neoplasia, and the second-leading cause of cancer-related death in women worldwide. Over 2 million cases are diagnosed annually, which represents the highest incidence of cancer in the world [1]. Also, obesity is a known risk factor for initiation, growth, invasion, and metastasis of breast cancer cells [2]. Metastasis is a complication of cancer in which neoplastic cells escape from the primary tumor, and develop secondary tumors in distant organs [3]. This process involves loss of cell-cell junctions and cell-extracellular matrix (ECM) interactions, acquisition of migratory capacity, matrix metalloproteases (MMPs) secretion, rupture of basement membrane, degradation of ECM and subsequently cell invasion [3]. Importantly, obese female patients present larger, advanced and more aggressive metastatic tumors to lymph nodes, than non-obese patients [4]. Obesity is characterized by an increase in adipokines production, which further enhance the predisposition of developing breast tumors [4]. Among these molecules, leptin is one of the most important adipokines involved in the development and progression of mammary tumors [5].

Leptin is a hormone with a molecular weight of ~16 kDa, encoded by the *OB* gene located on human chromosome 7 [6]. It is synthesized and secreted mainly by adipocytes, and in a smaller proportion, by the placenta, stomach, fibroblasts, skeletal muscle, and normal or tumorigenic epithelial mammary tissue [7]. One of the primary functions of leptin is the regulation of food intake and energy expenditure, acting primarily through the hypothalamus [8]. Leptin also regulates reproductive, immunological and metabolic functions [9]. Additionally, leptin is involved in the progression of breast cancer, through the activation of mitogenic, anti-apoptotic and metastatic pathways [2]. Leptin exerts these effects through the binding to the ObR receptor, activating various cellular signaling cascades such as JAK-STAT, MAPK and PI3K-Akt [7, 10]. Recent evidences showed that leptin levels in the plasma are higher in breast cancer patients compared with healthy individuals [2, 11, 12]. Furthermore, leptin and the ObR receptor are overexpressed in primary and metastatic mammary tumor tissues, suggesting an autocrine signaling mechanism developed by tumor cells [13, 14].

The focal adhesion kinase (FAK) participates in the formation of focal adhesions, and activates signaling pathways related to proliferation, survival, cell migration, and angiogenesis [15]. Classically, FAK is activated during the formation of focal adhesions, and it is mediated by the interaction between ECM with β-integrins, triggering conformational changes in these receptors [16]. This effect is followed by the autophosphorylation of FAK at Y397, which creates a high-affinity binding site for the Src-homology 2 (SH2) domain of the non-receptor tyrosine kinase Src, whose interaction promotes the autophosphorylation at Y418 of Src triggering its activation [17]. Active Src phosphorylates the Y576 and Y577 at the FAK kinase domain, leading to FAK maximum catalytic activity, and the formation of a transient FAK–Src signaling complex [15]. Considering this evidence, we hypothesized that leptin promotes FAK and Src activation, as well as metastasis-associated events such as cell migration, MMPs secretion and invasion.

In this study, we evaluated the role of leptin in the activation of FAK and Src kinases, and their roles in cell migration, MMPs secretion, and invasion in a cultured cell model of breast cancer. We utilized the breast cancer cell lines MCF7 and MDA-MB-231, and found that leptin induces FAK and Src activation. Furthermore, using a combination of inhibitors of these kinases we found a decrease in metastasis-associated events such as cell migration, MMPs secretion, and invasion in breast cancer cells. The data represented here contributes to the molecular characterization of the signaling events associated with leptin contributions to cell migration and invasion in breast cancer cell lines.

## MATERIALS AND METHODS

### Materials

Recombinant human leptin, FAK (PF-573228) and Src (PP2) inhibitors were obtained from Sigma-Aldrich (St Louis, MO, USA). Mouse anti-actin, rabbit anti-FAK and anti-Src antibodies were purchased from Santa Cruz Biotechnology (Santa Cruz, CA, USA). The phospho-specific antibody against FAK, rabbit anti-pY397, was obtained from Invitrogen (Carlsbad, CA, USA). The phospho-specific antibody against Src, rabbit anti-pY418, was obtained from MyBiosource (San Diego, CA, USA). Secondary HRP-conjugated antibodies were from Millipore (Billerica, MA, USA), and the secondary antibody anti-mouse/rabbit conjugated with Alexa Fluor 488 was from Invitrogen (Carlsbad, CA, USA). Phalloidin coupled to TRITC was purchased from Cytoskeleton (Denver, CO, USA).

### Cell culture

The breast cancer cell lines MCF7 and MDA-MB-231 (ATCC, Manassas, VA, USA) were cultured in DMEM/F12 media (50:50, V:V; Sigma-Aldrich, St Louis, MO) supplemented with 5% fetal bovine serum (FBS) and 1% antibiotics (penicillin G/Streptomycin, Gibco, Waltham, MA) in a humidified atmosphere containing 5% CO_2_ at 37 °C. For experimental purposes, cell cultures were serum-starved for 24 h before treatment with either FAK or Src inhibitors and/or leptin; and cell cultures were used between passages 3 and 15.

### Cell stimulation by leptin and kinase inhibitors

MCF7 and MDA-MB-231 cell cultures were seeded in 60 mm plates containing 4 ml of DMEM/F12. When the cell cultures reached confluence, they were washed with PBS, and then treated with FAK (5 μM) or Src (5 µM) inhibitors, and/or leptin for the times and concentrations indicated in the figure legends. Stimulation of confluent cells was terminated by removing the medium, and solubilizing the cells in 0.5 ml of ice-cold radioimmune precipitation assay (RIPA) buffer, containing 50 mM HEPES pH 7.4, 150 mM NaCl, 1 mM EGTA, 1 mM sodium orthovanadate, 100 mM NaF, 10 mM sodium pyrophosphate, 10% glycerol, 1% Triton X-100, 1% sodium deoxycholate, 1.5 mM MgCl_2_, 0.1% SDS and 1 mM phenylmethylsulfonyl fluoride (PMSF, Sigma-Aldrich).

### Western blot

Whole cell lysates (20 µg) were resolved on 10% SDS-polyacrylamide gels. Proteins were transferred to nitrocellulose membranes (Bio-Rad, Hercules, CA). The anti-Actin, anti-pY397, anti-FAK, anti-pY418 and anti-Src primary antibodies were incubated overnight at 4 °C, in agitation at a 1:1,000 dilution. Species specific secondary HRP-conjugated antibodies (1:5,000) were incubated for 2 h at room temperature. Membranes were developed using an enhanced chemiluminescence detection system from Bio-Rad.

### Immunofluorescence and confocal microscopy

MCF7 and MDA-MB-231 cells were seeded on glass coverslips, grown to 70% confluence, and stimulated either with or without leptin for 0, 15, 60 and 120 min. Cells were fixed for 5 min with 4% paraformaldehyde in PBS and permeabilized with 0.2% Triton-X100 in PBS at room temperature. For immunofluorescence assays (IF), the cells were blocked with 3% albumin in PBS for 1 h at room temperature. Then, the cells were incubated 2 h at room temperature with the rabbit anti-pY397 antibody (1:250 dilution), followed by a 2 h incubation at room temperature with an anti-rabbit conjugated to Alexa Fluor 488 secondary antibody (1:800 dilution). For F-actin staining, cells were incubated with TRITC-Phalloidin (1:500 dilution) for 30 min at room temperature. Cells were counterstained with 4′6-diamidino-2-phenylindole (DAPI), mounted with Fluoroshield/DAPI media (Sigma-Aldrich), and imaged with an Olympus BX43 microscope (Olympus Life Science, Center Valley, PA), using the 100X immersion objective.

### Wound healing assays

MCF7 and MDA-MB-231 cells were grown until confluence on 60 mm culture dishes supplemented with DMEM/F12 as described above. Cells were starved for 24 h in DMEM/F12 without FBS, and treated for 2 h with Cytosine ß-D-Arabinofuranoside (AraC) to inhibit cell proliferation during the experiment. After starvation, cells were scratch-wounded using a sterile 200 μl pipette tip, suspended cells were removed by washing with PBS twice, and the cultures were re-fed with DMEM/F12 in the presence or absence of the FAK and Src inhibitors, and/or leptin. The progress of cell migration into the wound was monitored at 0 and 48 h using an Olympus BX43 microscope with a 10X objective. The bottom of the plate was marked for reference, and the same field of the monolayers were photographed immediately after performing the wound (time = 0 h) and 48 h after treatments (time= 48 h), five images per plate were analyzed. The distance between the edges of the wound was measured at time 0 and 48 h, and the reported migrated distance corresponds to the difference between these two. The migration area was determined by measuring the total area of the wound using the ImageJ software and the MRI wound healing tool [18].

### Gelatin zymography

Cells were stimulated with leptin and the kinases inhibitors as described above, and the conditioned medium was collected, and concentrated using 30 kDa cut-off ultra-centrifugal filter units (Amicon, Merck-Millipore, MA). Protein was determined by Bradford [19], and 20 µg of protein from concentrated supernatant were assayed for proteolytic activity on native gelatin-substrate gels [20]. Briefly, samples were mixed with non-reducing buffer containing 2.5% SDS, 1% sucrose and 4 mg/ml phenol red, and separated in 8% acrylamide gels co-polymerized with 1 mg/ml gelatin, as previously described [21]. After electrophoresis at 72 V for 2.5 h, the gels were rinsed twice in 2.5% Triton X-100, and then incubated in 50 mM Tris HCl pH 7.4 and 5 mM CaCl_2_ assay buffer at 37 °C for 24 h. Gels were fixed and stained with 0.25% Coomassie Brilliant Blue G-250 in 10% acetic acid and 30% methanol. Proteolytic activity was detected as clear bands against the background stain of undigested substrate in the gel. Quantification was performed using ImageJ software [18].

### Cell Invasion assays

Matrigel invasion assays were performed following the Transwell chamber method [22], using 24 well plates containing inserts of 8 mm pore size (Corning Inc, Kennebunk, ME). Briefly, 30 µl of Matrigel (Corning) was added into the inserts and kept at 37 °C for 30 min to form a semisolid matrix. Untreated or treated cells with FAK or Src inhibitors were plated at 1×10^5^ cells per insert in serum-free medium on the top chamber. The lower chamber of the Transwell contained 600 µl DMEM supplemented or not with 100 ng/ml leptin. Cells were incubated for 48 h at 37 °C in a 5% CO_2_ atmosphere. Following incubation, cells and Matrigel on the upper surface of the Transwell membrane were gently removed with cotton swabs. Invading cells on the lower surface of the membrane were washed and fixed with methanol for 5 min, and stained with 0.1% crystal violet in PBS. Cell quantification was performed using a hemocytometer and an Olympus BX43 microscope with the 100X objective.

### Statistical analysis

Results are expressed as mean ± SD. Data was statistically analyzed using one-way ANOVA and the pairwise comparisons were performed using Newman-Keuls and Dunnett′s multiple comparison test. Statistical probability of P < 0.05 was considered significant.

## RESULTS

### Leptin induces the activation of FAK kinase in breast cancer cell lines

FAK activation is triggered by the autophosphorylation at Y397, which is an indicative of its catalytic activity [23, 24]. This process has been associated with cancer progression and metastasis-related events [24, 25]. First, we performed time-course assays to establish whether leptin induces FAK phosphorylation and activation in MDA-MB-231 and MCF7 breast cancer cell lines. We found that MDA-MB-231 cells treated with 100 ng/ml of leptin present a significant increase in FAK phosphorylation on Y397, after 10 min of treatment and peaked after 15 min compared to the non-treated control cells (Fig. 1A, C). Leptin also induces FAK activation in MCF7 cells, as Y397 phosphorylation is triggered after 15 min of treatment, reaching a maximum phosphorylation peak at 30 min (Fig. 1B, D). These data indicates that leptin-dependent FAK phosphorylation at Y397, and its consequent activation occurs shortly after the stimulus, but is not sustained for long periods in both breast cancer cell lines.

**Figure 1.**
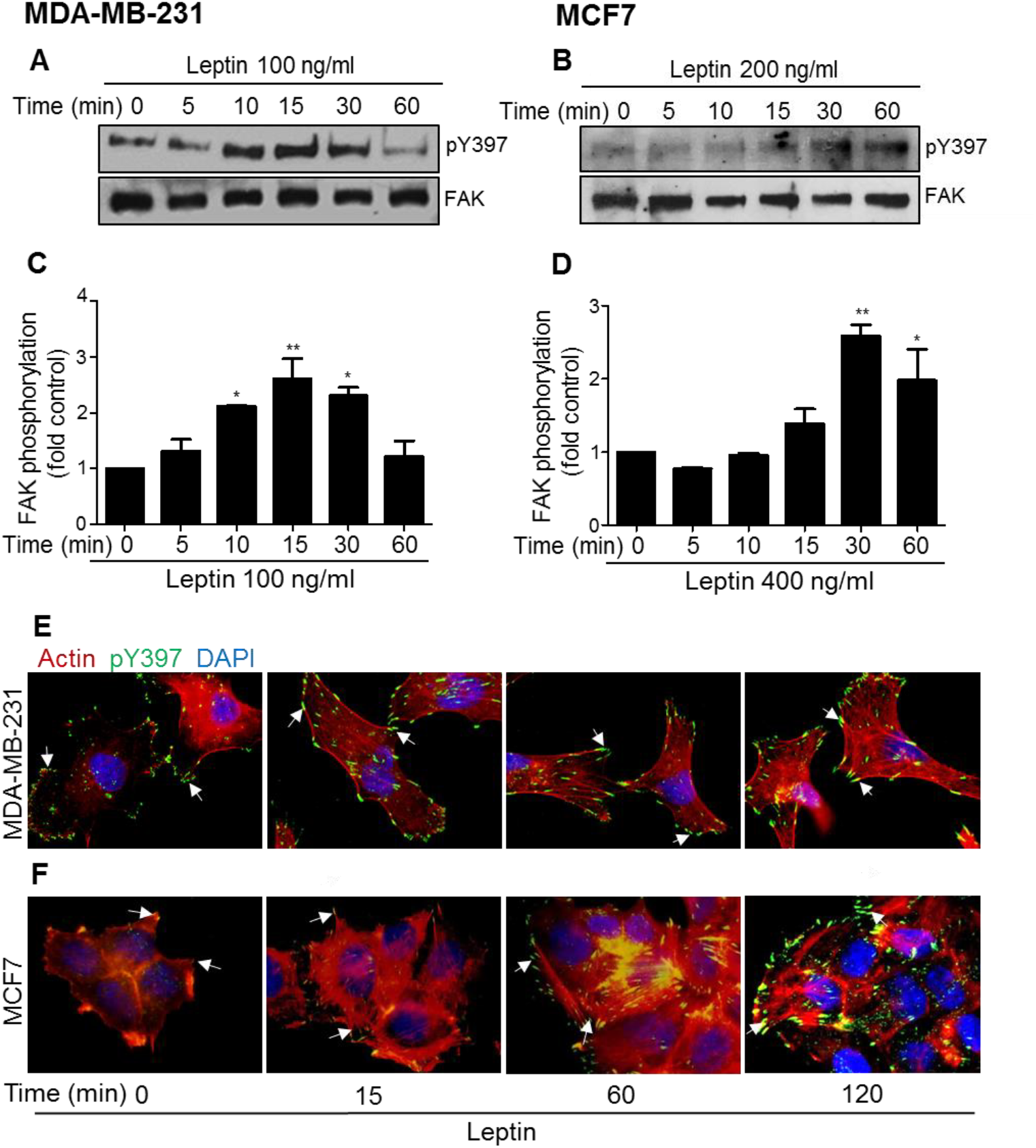
Activation and subcellular localization of FAK in leptin-stimulated breast cancer cell lines. MDA-MB-231 (A) and MCF7 (B) cells were treated with 100 and 200 ng/ml of leptin, respectively, and samples were collected at the indicated time points. Representative Western blots showing phosphorylated Y397 FAK and total FAK using specific antibodies. The graphs represent the densitometric and statistical analysis of the bands obtained by Western blot for MDA-MB-231 (C) and MCF7 (D); the values are the mean ± SD of at least three independent experiments, and are expressed as changes with respect to the control (unstimulated cells). Asterisks denote comparisons made to unstimulated cells. *P<0.05, **P<0.01 by one-way ANOVA (Dunnett test). Representative confocal microscopy images of MDA-MB-231 (E) and MCF7 (F) treated with or without leptin (100 and 200 ng/ml) at different time points. Anti-pY397 FAK staining is in green, TRITC-phalloidin is shown in red and DNA is stained with DAPI. Arrows indicate focal adhesions.

FAK activation is associated with the formation of focal adhesions and cell migration, as it contributes to the weakening of cell-cell adhesion and the cyclic assembly-disassembly of focal adhesions [26]. Therefore, we analyzed the effect of leptin in the subcellular localization of phosphorylated FAK at Y397, and whether its activation would promote the formation of stress fibers in MDA-MB-231 and MCF7 cells. Cells were incubated with leptin for 15, 30 and 60 min and formation of stress fibers was monitored by confocal microscopy. Optical planes taken in proximity of the cells with the substrate showed that under normal conditions, the MDA-MB-231 cells expressed activated FAK at the periphery of the cells, which increased upon leptin stimulus (Fig. 1E, in green). In these cells, the cytosolic actin filaments were also abundant, and also increased upon the treatment of leptin (Fig. 1E, in red). Interestingly, the effect of leptin in FAK activation, and formation of stress fibers was more evident in MCF7 cells. Confocal microscopy analyses of MCF7 cells showed low levels of activated FAK at the periphery of untreated cells (Fig. 1F, in green), which increased throughout the treatment with leptin. Importantly, these cells also had increased formation of stress fibers upon leptin stimulation (Fig. 1F, in red). These results suggest that leptin promotes the phosphorylation of FAK at Y397, and induces the formation of focal adhesions enriched with stress fibers. This hypothesis is in agreement with previous reports of FAK associated with cell migration in MDA-MB-231 and MDA-MB-468 in breast cancer cell lines [27], and with the effect of Y397 autophosphorylation in cell migration, invasion, and proliferation of gastric carcinomas [25].

### Leptin promotes cell migration in a FAK-dependent manner in MDA-MB-231 and MCF7 breast cancer cell lines

The increased phosphorylation of FAK at focal adhesion points of MDA-MB-231 and MCF7 cells is consistent with the requirement of this kinase for the cycling of focal adhesions during cell migration [28]. These observations prompted us to investigate whether FAK was part of the effect yielded by leptin in the migration of these cancer cell lines. Figure 2 shows representative light microscopy images of MDA-MB-231 and MCF7 cells subjected to wound healing assays. Time 0 h represents images of the wounds produced at the moment of scratching, while 48 h depicts the migration of both cell lines after performing the wound. The morphological changes observed at the front edge of the wound of both cell lines treated with leptin rendered a mesenchymal phenotype; while the cells treated with the specific FAK inhibitor, PF-573228, and leptin exhibited an epithelial phenotype (Fig. 2A, B). Both cell lines treated with leptin migrated significantly faster than non-treated control cells (Fig. 2C, D). However, in the presence of leptin, the MDA-MB-231 cell line (Fig. 2A, B) presented enhanced migration capabilities, compared to MCF7 cells (Fig 2A, C). Moreover, the MDA-MB-231 cells treated with the FAK inhibitor, also presented a larger decrease in leptin-dependent induced cell migration compared to MCF7, and to the leptin treated cells (Fig. 2). Similar trend in the phenotype was observed in MCF7 cells treated with the FAK inhibitor, however was reduced compared to MDA-MB-231 (Fig 2A, C). MCF7 cells presented approximately a 30% decrease in the leptin-induced migration properties when treated with the chemical inhibitor of FAK, compared to the leptin treated cells (Fig. 2A, C). These results suggest that leptin-induced migration is dependent on FAK kinase-mediated signaling pathways in both cell lines.

**Figure 2.**
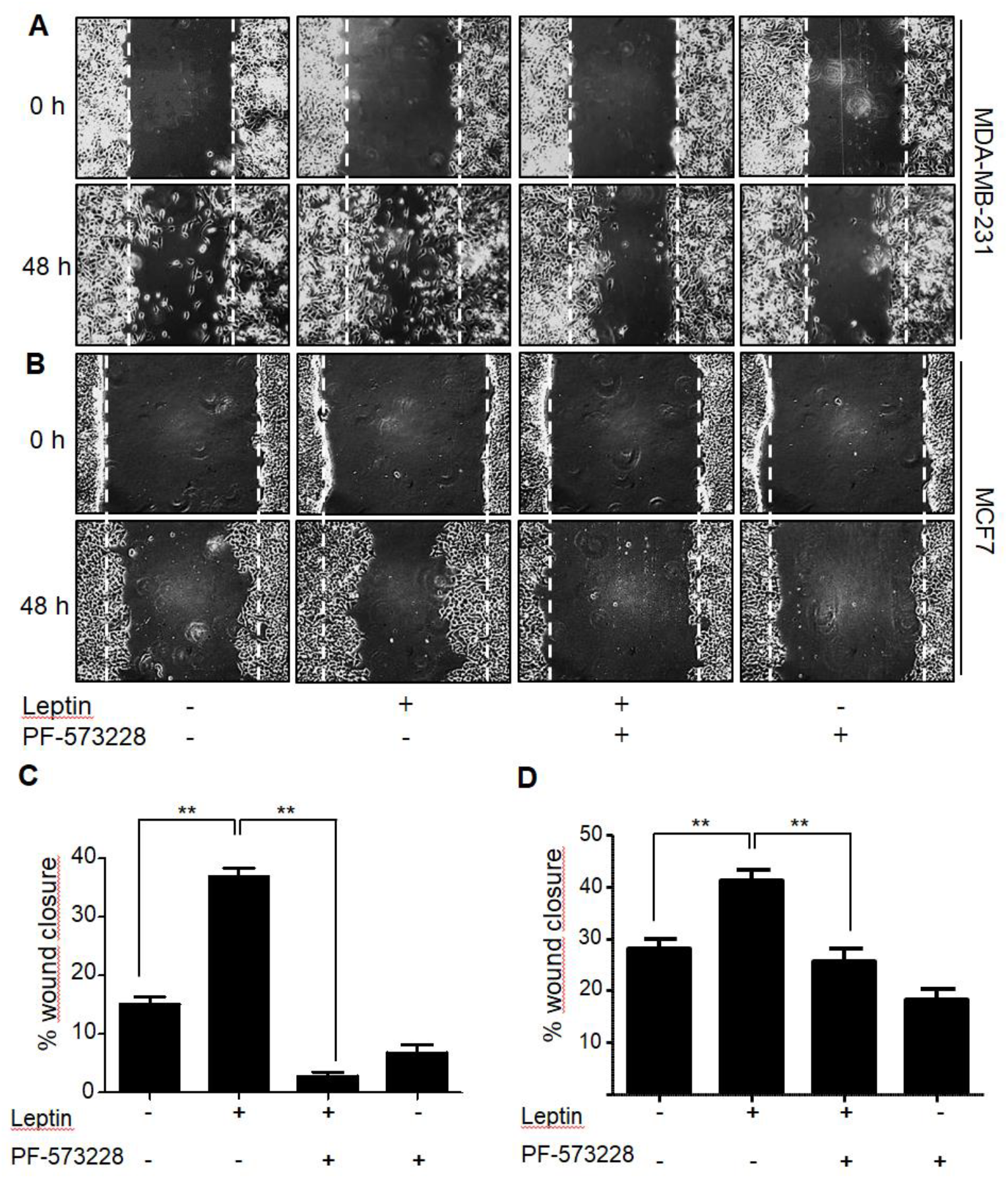
Leptin promotes cell migration in a FAK-dependent manner of MDA-MB-231 and MCF7 cell lines. Representative light microscopy images of wounding assay in MDA-MB-231 (A) and MCF7 (B) breast cancer cell lines. Confluent monolayers of MDA-MB-231 and MCF7 cells were treated for 2 h with AraC to inhibit proliferation, and then pre-treated with 5 µM of the FAK inhibitor, PF-57322, for 30 min. Wound healing assay was performed by scratching the monolayer using a sterile pipette tip. Cells were washed and re-fed with DMEM supplemented with or without leptin. The progress of cell migration into the wound was registered at 0 (upper panel) and 48 h (lower panel). Quantification of MDA-MB-231 (C) and MCF7 (D) percentage of wound closure for each condition and represents the mean of three independent experiments ± SD. The Newman-Keuls test compares all experimental conditions to determine statistically significant differences; **P <0.01 per one way ANOVA; **P < 0.01 by one-way ANOVA.

### Leptin increased metalloproteases secretion and activation in breast cancer cells

During migration, the cells need to modify the composition of the ECM by secreting MMPs. MMPs are secreted in response to growth factors, hormones, and cytokines [29]. To be active, the MMPs bind to divalent cations, such as Zn^2+^ [30]. MMPs are required for cell migration, wound healing, tissue remodeling, and angiogenesis among other processes, features by which MMPs are highly relevant in diseases such as arthritis and cancer [31]. MMPs secretion, in particular MMP-2 and MMP-9, has been associated with invasive and metastatic properties of cancers cells [32].

Therefore, we asked whether leptin induces the secretion and activation of MMP-2 and MMP-9. To test this hypothesis, MDA-MB-231 (Fig. 3A-C) and MCF7 cells (Fig. 3G-I) were stimulated with increasing concentrations of leptin for 24 h. The supernatants were collected, concentrated, and analyzed by gelatin zymograms to determine the activity of both MMPs. Figure 3 shows that leptin induced a gradual increase in MMP-2 and MMP-9 secretion and activation, which correlated with the amount of leptin supplemented into the culture media. In both cancer cell lines, a significant difference in the activity of MMP-2 and MMP-9 was observed. In the case of MDA-MB-231 cells, the major change in activation of MMP-2 occurred at 100 ng/ml of leptin, while the activity of MMP-9 was significantly higher at 50 ng/ml (Fig. 3D-F) compared with the non-treated controls. In the case of MCF7 cells, the activity of MMP-2 was significantly higher at 200 ng/ml of leptin, and for MMP-9 was at 100 ng/ml of the hormone (Fig. 3G-I). In both cases, the maximum peak of secretion and activation was achieved at 400 ng/ml (Fig. 3A-H).

**Figure 3.**
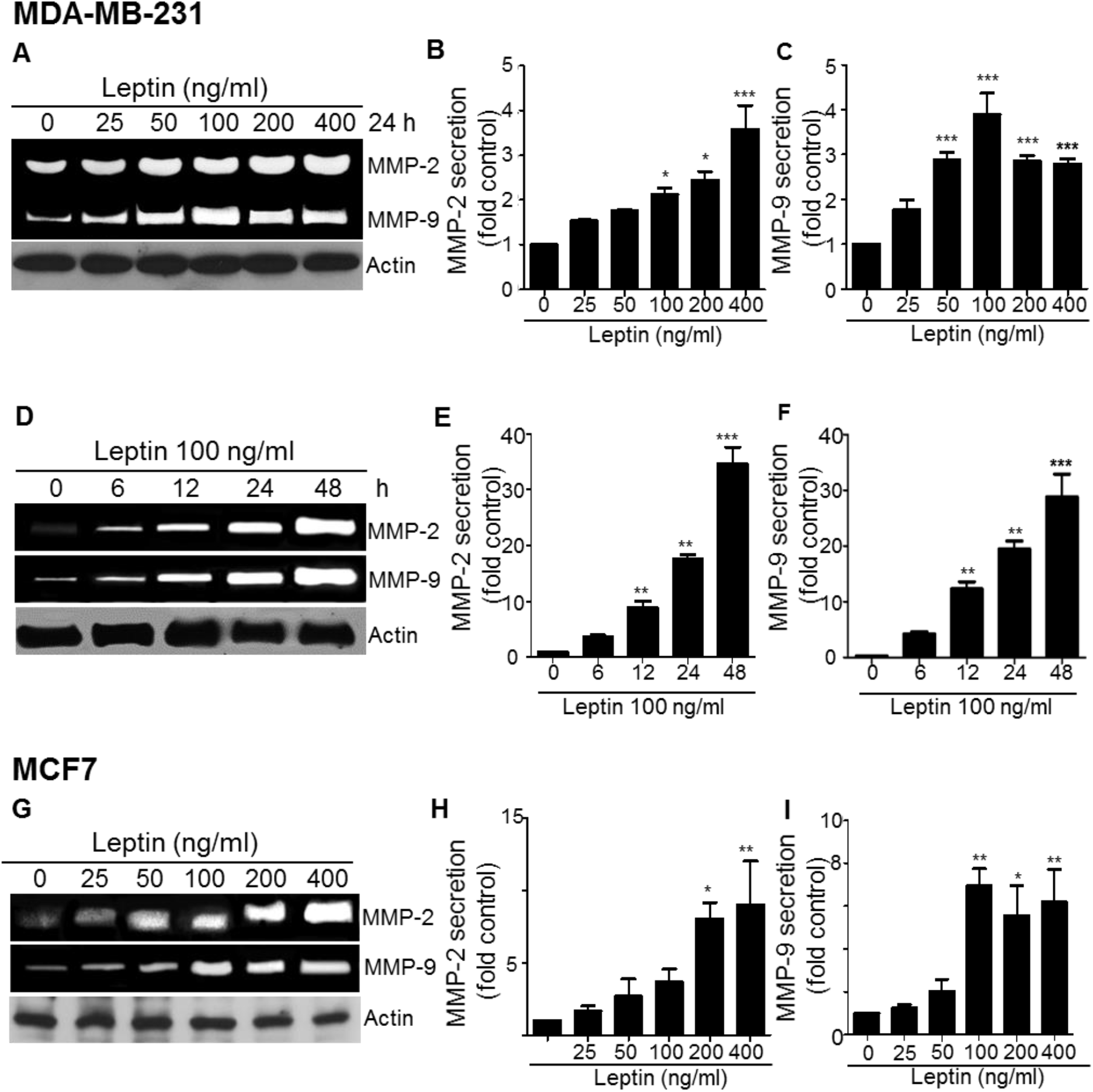
Leptin promotes the secretion of MMP-2 and MMP-9 in the MDA-MB-231 and MCF7 cancer cell lines. (A) Representative zymogram of the gelatinase activity of secreted MMP-2 and MMP-9 obtained from supernatants of MDA-MB-231 cells stimulated with increasing concentrations of leptin for 24 h. Quantification of the gelatinase activity of both secreted metalloproteases MMP-2 (B), MMP-9 (C) under each experimental condition. (D) Representative zymogram of the gelatinase activity of secreted MMP-2 and MMP-9 obtained from supernatants of MCF7 cells stimulated with increasing concentrations of leptin for 24 h. Quantification of the gelatinase activity of both secreted metalloproteases MMP-2 (E), MMP-9 (F) under each experimental condition. (G) Representative zymogram of the gelatinase activity of secreted MMP-2 and MMP-9 obtained from supernatants of MDA-MB-231 cells chronically stimulated with 100 ng/ml of leptin. Supernatant samples were collected at the indicated time points. Actin was used as loading control in all cases. Data represents the mean of three independent biological experiments ± SD and it is expressed as fold increase over the un-treated control. Statistical significance was established at *P<0.05, **P<0.01, ***P<0.001 by Dunnett test.

To determine the time frame for secretion and activation of the MMPs induced by leptin, we collected supernatants of MDA-MB-231 cells stimulated with 100 ng/ml of leptin for 6, 12, 24 and 48 h and assayed for gelatinase activity. Figure 3 G-I show that leptin induces an increase in both, MMP-2 and MMP-9 secretion after 6 h, reaching a maximum activity and secretion peak at 48 h. Together, our data suggest that MMP-2 and MMP-9, two relevant proteins for cell migration and metastatic processes, may be induced and activated by the exposure of chronic and increasing doses of leptin. These processes may have a direct correlation with the enhanced mobility capabilities of these cells observed in our wound healing assays (Fig. 2).

### Leptin promotes invasion in a FAK-dependent manner in MDA-MB-231 cells

MMPs secretion is directly related with invasive capacity of tumor cells [33], and might be one of the multiple mechanisms by which FAK positively regulates cell migration [23]. If MMP-2 and MMP-9 participate in leptin-dependent migration via FAK activation, their secretion must be controlled by similar signaling pathway that controls migration of cancer cells. To test this hypothesis, we used the specific FAK inhibitor, PF-573228, and evaluated the effect of inhibiting this signaling pathway on the leptin-induced secretion and activation of MMP-2 and MMP-9. MDA-MB-231 cells were pre-treated with the FAK inhibitor. Then the cells were stimulated with 50 ng/ml of leptin, the supernatants were collected, and concentrated for zymogram analyses. We found that pre-treatment with PF-573228 significantly inhibited MMP-2 (Fig. 4A, B) and MMP-9 (Fig. 4A, C) secretion and activation, when compared to the leptin treated cells.

**Figure 4.**
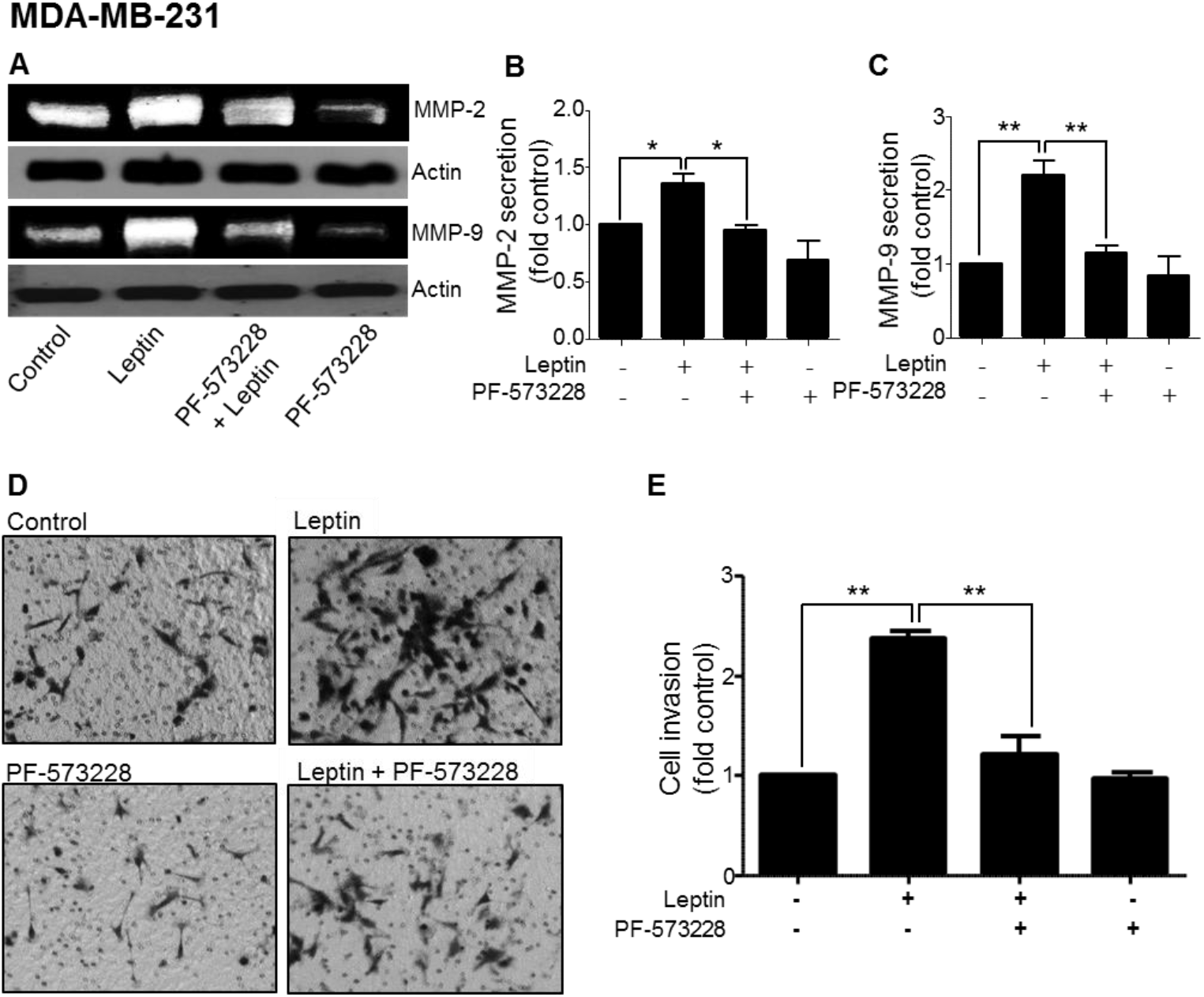
Leptin promotes FAK activation, secretion of metalloproteases and increases the invasive phenotype of MDA-MB-231 cell line. MDA-MB-231 cells were pre-treated for 30 min with 5 μM of the FAK-specific inhibitor, PF-573228. Then, the cells were stimulated without or with 100 ng/ml leptin for 24 h. (A) Representative zymogram of the gelatinase activity of MMP-2 and MMP-9 obtained from MDA-MB-231 cells supernatants, actin was used as loading control. Quantification of the activity of secreted MMP-2 (B) and MMP-9 (C) for each experimental condition, and it is expressed as fold increase over the un-treated control. Statistical significance was established at *P < 0.05, **P<0.01, ***P<0.001 by Dunnett test. (D) Representative light microscopy images of invasion assays for MDA-MB-231 cells. The cells were pre-treated with 5 μM PF-573228 and seeded at 1×105 cells in Matrigel-coated chambers in the presence or absence of 100 ng/ml leptin for 48 h. (E) Cell invasion quantification represents the number of invading cells for each experimental condition. Data represents the mean ± SD, and it is expressed as fold increase over the un-treated control. Statistical significance was established at *P<0.05, **P<0.01, by one-way ANOVA (Newman-Keuls test).

Further, we tested the role of FAK in the invasiveness capabilities of MDA-MB-231 cells stimulated with leptin. Cell invasion assay allows the evaluation of the intrusive potential of tumor cells through extracellular matrices, which is directly related to processes related to tumor progression such as metastasis [34]. Therefore, we sought to investigate the capabilities of the cells to penetrate a barrier consisting of components of the basement membrane in response to leptin and the FAK inhibition. To this end, the cells were seeded in Matrigel-coated chambers in the presence or absence of 100 ng/ml leptin for 48 h. Figure 4D shows representative light microscopy images of MDA-MB-231 cultured in the presence and absence of leptin, where the hormone promoted an increase in cell invasion, as shown by the presence of darker cells which denotes cell invasion. Also, we observed that invasiveness is a FAK-dependent process, as the cells treated with 5 μM of PF-573228, showed a decrease in invasiveness properties, compared to cells treated with leptin (Fig. 4D, E). These data is consistent with the contribution of FAK to the invasion properties of the triple-negative MDA-MB-231 and BT549 cancer cells [27, 35].

### Leptin promotes invasion in a FAK/Src-dependent manner in MDA-MB-231 cells

The classic mechanism for FAK activation involves autophosphorylation at the Y397, which creates a high-affinity binding site for Src [23]. The interaction between Src and FAK leads to the phosphorylation, and activation of Src at Y419, which further result in the phosphorylation of FAK at Y576 and Y577 [23, 36]. Src is a tyrosine kinase involved in cell migration and invasion in different triple negative cancer cell lines [37].

Therefore, we asked whether leptin also played a role in the activation of Src, using the highly invasive triple negative cell line MDA-MB-231 as an *in vitro* model. Figure 5A and B shows that leptin induces Src phosphorylation at Y418 time-dependent manner, beginning 10 min after stimulation and is sustained for at least 1 h.

**Figure 5.**
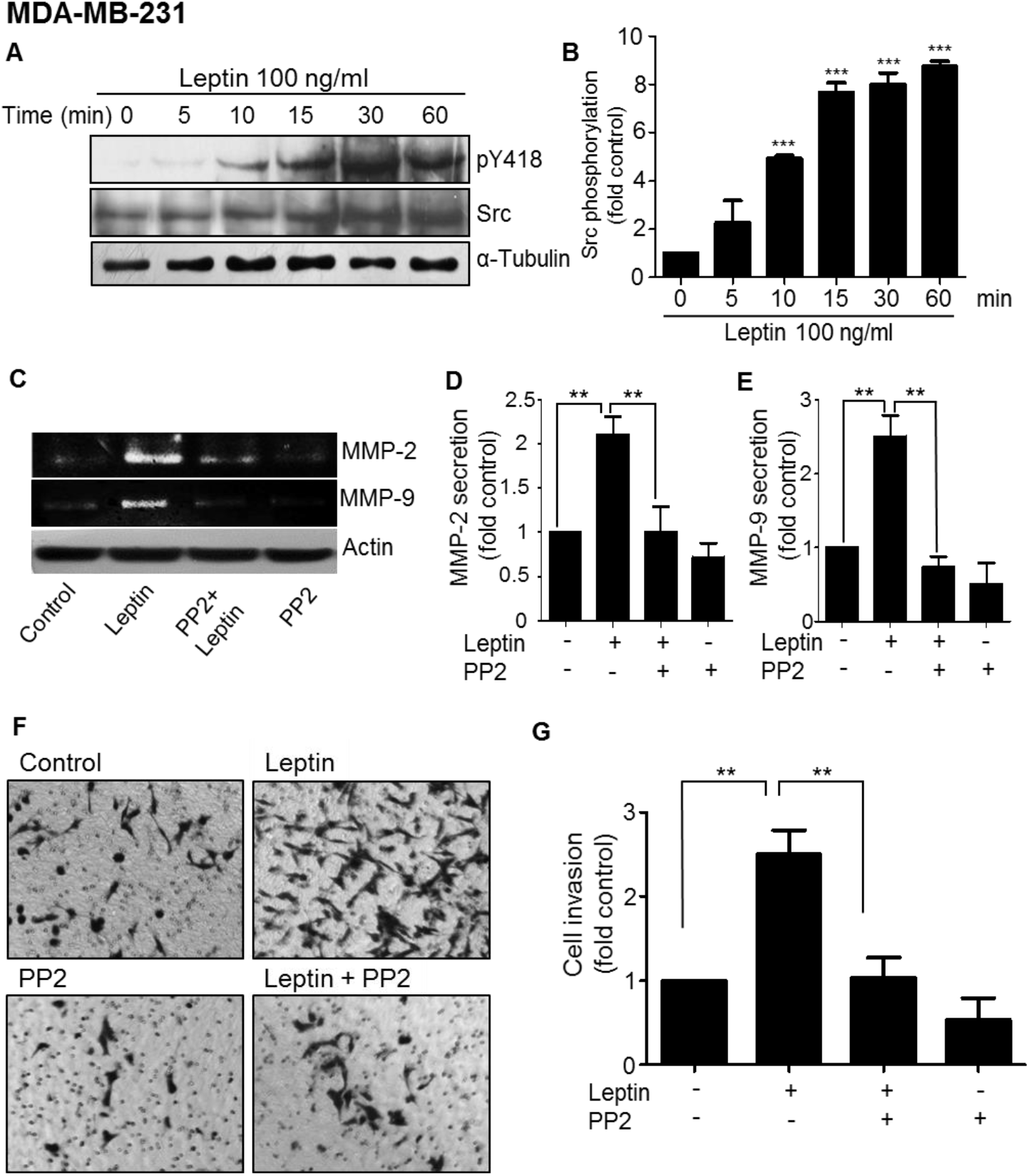
Leptin-dependent Src activation, leads to secretion, activation of MMP-2 and MMP-9, and invasion in MDA-MB-231 cells. (A) Representative Western blot of MDA-MB-231 cells treated with 100 ng/ml leptin at different time points. Phosphorylated Src was detected using an anti-pY418 as well as total Src. (B) Densitometric and statistical analysis of three independent Western blot analyses of phosphorylated Src at pY418. Data represents the mean ± SD, and are expressed as changes with respect to the un-treated control. Statistical significance was established at *P<0.05, **P<0.01 by Dunnett test. (C) Representative zymogram MMP-2 and MMP-9 obtained from supernatants of MDA-MB-231 cells treated with 5 μM of the Src-specific inhibitor, PP2, and supplemented with or without 100 ng/ml leptin for 24 h. Quantification of the activity of secreted MMP-2 (D) and MMP-9 (E) for each experimental condition, and it is expressed as fold increase over the un-treated control. Statistical significance was established at *P<0.05, **P<0.01, by one-way ANOVA (Newman-Keuls test). (F) Representative light microscopy images of invasion assays for MDA-MB-231 cells. The cells were pre-treated with 5 μM of PP2 and seeded at 1×105 cells in Matrigel-coated chambers in the presence or absence of 100 ng/ml leptin for 48 h. (G) Cell invasion quantification represents the number of invading cells for each experimental condition. Data represents the mean ± SD, and it is expressed as fold increase over the un-treated control. Statistical significance was established at *P<0.05, **P<0.01, by one-way ANOVA (Newman-Keuls test).

Finally, we evaluated the role of Src in the secretion and activation of metalloproteases in MDA-MB-231 cells. We observed that leptin promotes the secretion of MMP-2 (Fig. 5C, D), and MMP-9 (Fig. 5C, E) through the activation of Src, since incubation of the cells with the specific inhibitor of this kinase, PP2, prevented this process. This phenotype was accompanied by a decrease in the leptin-induced invasive capabilities of the MDA-MB-231 cell line treated with PP2 (Fig. 5F, G), supporting the role for Src in cell migration and invasion of cancer cells.

Taken together, our data suggest that leptin induces a FAK/Src-dependent activation and secretion of MMP-2 and MMP-9, which may contribute to the cell migration and invasion phenotypes observed in *in vitro* models of mammary cancer cells. This leptin-dependent phenotype associated with the activation of FAK and Src is consistent with a more aggressive phenotype of the tumorigenic and metastatic cancer cells.

## Discussion

Obesity is considered one of the risk factors associated with the development and progression of breast cancer [5, 38]. Adipose tissue is characterized by an increased synthesis of different adipokines such as leptin [14]. Leptin regulates several physiological functions such as food intake and energy expenditure [8]. However, *in vitro* studies demonstrated that leptin is also an inducer of the epithelial-mesenchymal transition in breast epithelial cells [39–41], and that it is associated with tumor progression in breast cancer [7]. Leptin is also secreted by adipocytes in the tumor microenvironment, which induces cell migration in breast cancer cell lines [42]. Cell migration is a key step in metastasis of tumor cells, and occurs via two possible mechanisms for mobility: 1) amoeboid, 2) mesenchymal patterns [43]. While the amoeboid-type of migration has been reported to be independent of integrins and proteases [44], the mesenchymal migration appears dependent on integrins, proteases and signaling molecules such as FAK [28]. FAK is essential for the formation of focal adhesions and cell migration [23].

In this study, we used the non-invasive MCF7 breast cancer cell line, and the triple negative highly invasive MDA-MB-231 to investigate the signaling pathways underlying cell migration and invasion. Our results showed that leptin promotes the phosphorylation and consequent activation of FAK at Y397, in both cell lines; however, the phenotypes observed in the triple negative MDA-MB-231 were significantly enhanced, compared to the MCF7 cells. We observed that in both cell lines, leptin induces the activation of FAK at 10 min of stimulation. Data from the colon cancer cell line SW480 reported by Ratke *et al.* 2010, showed that leptin induces FAK activation in a time-specific manner with a maximal activation at 15 min [45], which is consistent with our studies. Although the exact mechanism of leptin-induced FAK activation has not been described, experimental evidence showed that its canonical activation pathway is through the autophosphorylation of Y397 [46], which is in agreement with our observations. This process is a response to the binding of integrins to ECM during the formation of focal adhesions [15]. However, this might not be the only pathway for FAK activation. It has been reported that receptors with tyrosine kinase activity may promote FAK activation in NIH 3T3 fibroblastic cell lines [47]. Consistently, in our study FAK phosphorylated at Y397 is localized primarily in focal adhesions bound to stress fibers in both breast cancer cell lines, which suggest that this subcellular localization promotes cell migration.

Considering that leptin induces activation of FAK in these breast cancer cell lines, we evaluated the effect of FAK on leptin-induced cell migration by using the chemical inhibitor PF-573228. We found that leptin promotes migration of breast cancer cells in a FAK kinase-dependent manner. Focal adhesions are specialized structures where integrin receptors interact with the ECM, and with the actin cytoskeleton to promote cell migration [48]. In addition to regulating the formation of focal adhesions, FAK kinase signaling also affects the remodeling of the actin cytoskeleton by inducing the activation of small GTPases such as Rho, Rac and Cdc42 [49]. These are associated with the reorganization of the actin cytoskeleton by generating structures involved in cell migration, such as lamellipodia, filopodia, and stress fibers [50].

An important feature of tumor progression in cancer cells is the increased secretion and activation of metalloproteases [51]. We focused our studies in MMP-2 and MMP-9 because they are related to invasive and metastatic processes [52]. The biological relevance of the expression, secretion, and activation of MMPs, is their relation with a highly invasive capacity of tumor cells [53]. In particular, MMP-2 and MMP-9 degrade type IV collagen and promote the rupture of basal membranes in colorectal, prostate, lung and breast cancer [54, 55]. Here, we showed that in two breast cancer *in vitro* cellular models, leptin induces an increase in MMPs secretion and activation in a time and dose response-dependent manner. These findings are in agreement with evidence collected from serum samples from breast cancer patients, where high levels of MMP-2 and MMP-9 has been directly associated with metastasis, and further provide evidence of the signaling mechanisms [56].

MMP-2 and MMP-9 expression is connected with different phases of metastasis, such as the pre-metastatic niche formation, formation of new blood vessels and local invasion [57]. We showed that leptin promotes cell invasion of the triple negative MDA-MB-231 cell line in a FAK-dependent pathway. Our results suggest that leptin exposure promotes FAK activation, and in consequence downstream signaling cascades leading to the secretion and activation of both metalloproteases. FAK-dependent induction of MMPs is not limited to cancer; in tenocytes the mechano-growth factor promotes MMP-2 secretion and cell invasion in a manner dependent on FAK and ERK kinases [58]. MMP-9-dependent degradation of fibronectin also stimulates cell migration, and invasion of MCF-7 cells in a FAK/Src-dependent manner [59]. The relevance of FAK/Src in cell migration and invasive processes extends to other cellular systems. For instance, the cell adhesion protein Spondin 1, involved in the attachment of sensory neuron cells and growth of neurites, also promotes cell migration and invasion through a FAK/Src-dependent pathway in human osteosarcoma cell lines [60]. Further, Src is also associated with cell permeability, EMT, migration, invasion and metastasis of tumor cells [61]. In this work, we evaluated the involvement of Src kinase in the secretion of metalloproteases and leptin-induced cell invasion. We found that leptin promotes the activation of Src which led to the secretion of metalloproteases and invasion events. Together, these data suggest that leptin is associated with the establishment of a more aggressive phenotype of the tumor cells promoting local invasion and eventually metastasis of tumor cells (Fig. 6).

**Figure 6.**
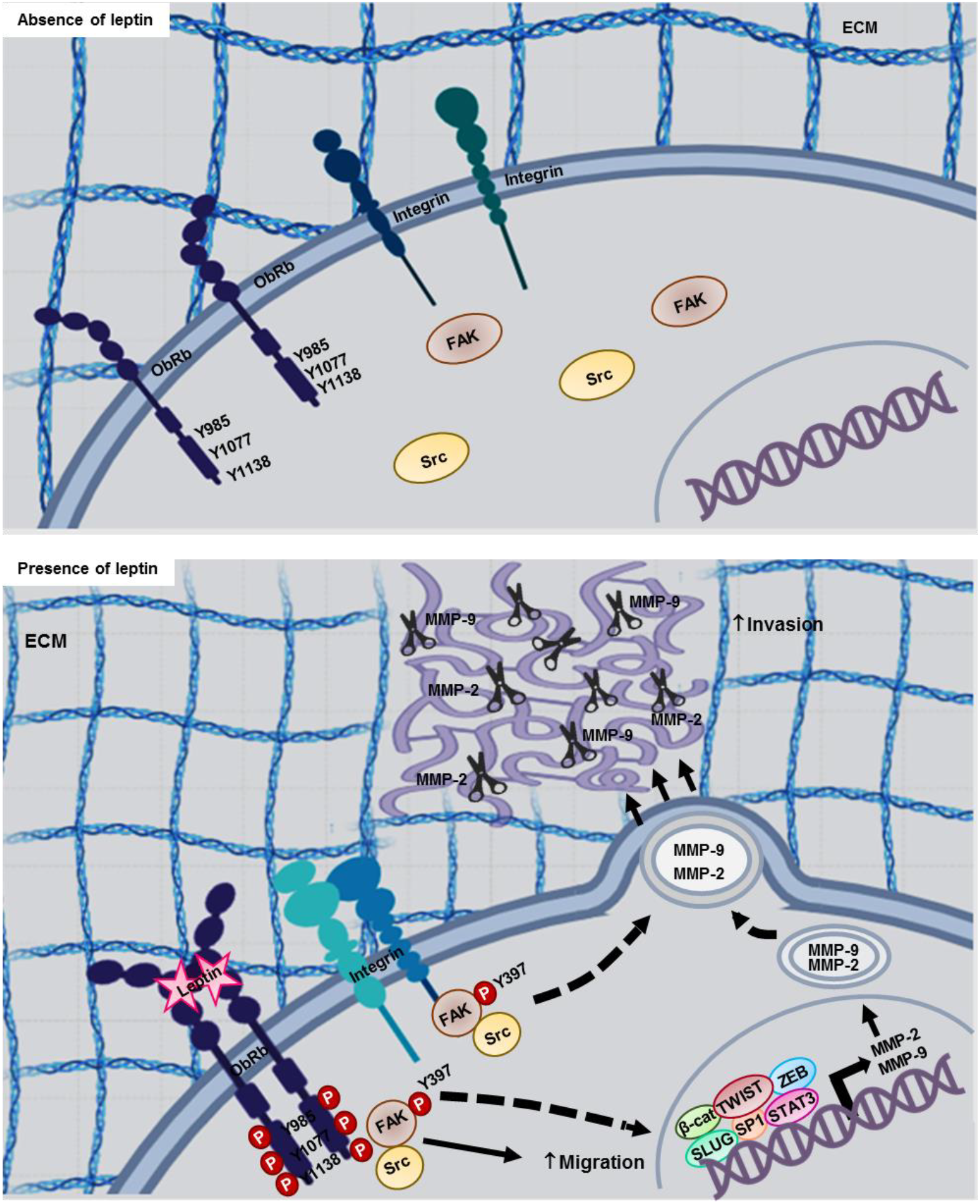
Representative model of the MMP-2 and MMP-9 secretion induced by leptin in a cultured cellular model for breast cancer. Leptin binding to the ObRb receptor activates FAK and Src kinases, and signaling cascades that promotes the secretion of MMPs, cell migration and invasion in breast cancer cells.

## Conclusion

Taken together, our results demonstrate that leptin promotes cell migration, MMPs secretion and invasion in a FAK and Src-dependent pathway. These events partially explain the association between obesity and the development and progression of breast cancer and suggest that kinases FAK and Src are central molecules, regulating events that favor the metastasis of tumor cells stimulated with leptin, promoting changes to a more aggressive phenotype in breast cancer cells.

## Acknowledgements

We thank Travis Ashworth for the style corrections of this manuscript.

